# Enrichment of long DNA fragments from mixed samples for Nanopore sequencing

**DOI:** 10.1101/048850

**Authors:** SE Eckert, JZ-M Chan, Darren Houniet, the PATHSEEK consortium, J Breuer, G Speight

## Abstract

Whole-genome sequencing of pathogenic organisms directly from clinical samples combines detection and genotyping in one step. This can speed up diagnosis, especially for slow-growing organisms like *Mycobacterium tuberculosis (Mtb)*, which need considerable time to grow in subculture, and can provide vital information for effective personalised treatment. Within the PATHSEEK project, we have developed a bait-capture approach to selectively enrich DNA/RNA from specific bacterial and viral pathogens present in clinical samples. Here, we present a variation of the method that allows enrichment of large fragments of target DNA for sequencing on an Oxford Nanopore MinION^TM^ sequencer. We enriched and sequenced cDNA from Influenza A (FluA), genomic DNA (gDNA) from human cytomegalovirus (CMV) and from two strains of *Mtb*, and present an evaluation of the method together with analysis of the sequencing results from a MinION^TM^ and an Illumina MiSeq sequencer. While unenriched FluA and CMV samples had no reads matching the target organism due to the high background of DNA from host cell lines, enriched samples had 56.7% and 90.9% on-target reads respectively for the best quality Nanopore reads.

## Introduction

Amidst increased occurrence of extensive or totally drug-resistant pathogens and antibiotic overuse^1^, high-throughput sequencing methods, particularly those which can be readily applied in a clinical setting, are being used to aid and refine diagnosis in a timely fashion^2^. Data from wholegenome sequencing provides a wealth of information such as identification of resistance markers carried by the infecting agent(s), allowing for rapid, targeted and personalised treatment. However, DNA extracted from clinical samples consists of a mixture of low amounts of pathogen genetic material and overwhelming quantities of human and commensal DNA. Previous studies have shown that it is possible to bypass the traditional culture-based diagnosis and obtain informative sequence data from metagenomic samples, but the throughput is low and the method prohibitively costly^3^. The EU-funded FP7 PATHSEEK project (http://www.pathseek.eu/) has developed a disruptive diagnostic platform to sequence bacterial and viral pathogens directly from clinical samples^4, 5^. This enrichment approach employs custom baits to capture genomic material from the target pathogens, thereby removing the human and commensal DNA, and allowing greater throughput of samples on Illumina sequencers. However, as this method is optimised for short-read sequencers such as the Illumina MiSeq and the Ion PGM, it is very difficult to resolve highly repetitive regions such as those found in cytomegalovirus^6^, or provide evidence of recombination events such as those seen in Chlamydia trachomatis. Members of the PATHSEEK consortium joined the Oxford Nanopore Technologies (ONT, Oxford, UK) MinION^TM^ access program to assess the suitability of its long-read platform, the MinION^TM^, for the targeted enrichment method.

Here, we present an adaption to the PATHSEEK method – enrichment of DNA fragments of between 1 and 15 kb for sequencing on long-range platforms. We compare sequence data from unenriched and enriched cultured FluA and CMV samples, run on the MinION^TM^ and Illumina (San Diego, CA, USA) MiSeq platforms. We also mixed cultured *Mtb* gDNA from two different strains with human DNA to assess the efficiency of enrichment by hybridisation for longer bacterial DNA fragments. Long genomic fragments were readily purified from a background of the cell line used for producing the viruses, or, in case of *Mtb*, admixed human DNA.

## Materials and Methods

### Samples

Mycobacterial gDNA from strains H37Rv and the extensively drug-resistant clinical strain *Mtb*C were kind gifts from A. Brown, L. J Schreuder, T Parish (Barts and The London School of Medicine and Dentistry, Queen Mary University of London, London, UK), P. Butcher and J. Dhillon^7^ (Institute of Infection and Immunity, St. George’s Hospital, University of London, UK). To simulate clinical samples, *Mtb* DNA was mixed with human gDNA (Male, #G1471, Promega, Madison WI, USA), to 10% (450 ng human DNA, 50 ng *Mtb* DNA) or 90% (450 ng *Mtb*, 50 ng human) prior to processing.

RNA from Influenza strain A A/PR/8/34 #0111041v H1N1, grown in MDCK Cocker Spaniel kidney cell line, was obtained from the Public Health England Culture Collection (Porton Down, UK), and reverse transcribed with NEBNext RNA First Strand Synthesis Module #E7525 and NEBNext mRNA Second Strand Synthesis Module #E6111 (New England Biolabs, Hitchin, UK) according to the manufacturer’s instructions. The sample was cleaned up with DNA Clean & Concentrator columns (#D4013, Zymo Research, Irvine CA, USA).

DNA from CMV strain Merlin grown in fibroblast cell culture was subjected to PreCR (#M0309, New England Biolabs, Ipswich MA, USA) enzymatic repair according to the manufacturer’s recommendations after shearing. It was cleaned up with 100 μl (1 volume) Agencourt AMPure XP beads (#A63880, Beckman Coulter High Wycombe, UK), according to the manufacturer’s instructions, and eluted in 25 μl H_2_O.

### Sample preparation and long-fragment hybridisation

CMV and *Mtb* Samples (500 ng) were diluted in TE to an end volume of 80 µl, and sheared in Covaris g-TUBEs (#520079, Covaris, Woburn MA, USA) with two passages at 7200 rpm/4200 g for 1 min in a desktop centrifuge (#5242, Eppendorf, Hamburg, Germany). FluA samples were not sheared as the cDNA fragments, derived from eight segments of negative-sense RNA of approximately 700 – 2300 nucleotides (nt), were size-compatible with Nanopore sequencing.

Concentrations and fragment sizes were determined with a Qubit fluorometer (dsDNA BR Assay Kit #Q32850, Life Technologies Ltd, Paisley, UK), and Agilent Tape Station (Genomic DNA ScreenTape #5067-5365 and Genomic DNA Reagents #5067-5366, Agilent, Santa Clara CA, USA) according to manufacturers’ instructions.

PATHSEEK custom baits for the target organisms FluA, CMV and *Mtb* were designed using an inhouse Perl script developed by the PATHSEEK consortium, using a database of 4968 H1N1 and 2966 H3N2 FluA genomes, 115 partial and complete CMV genomes and the *Mtb* strain H37Rv reference genome (AL123456.3) respectively. Sheared gDNA (CMV, *Mtb*) and cDNA (FluA) samples (500 ng) were hybridised and captured using 2 μl FluA and CMV baits, or 5 μl for *Mtb* baits per reaction. Hybridisation and washing were performed using a modified version of the SureSelect^XT^ Target Enrichment for Illumina Paired-End Multiplexed Sequencing as described previously^4, 5^. The workflow used in this study is outlined in Figure 1. Samples (30 μl) were then heated to 95°C for 3 min, and cooled to 35°C (ramp: 4°C/min) to release the target fragments from the baits and streptavidin beads.

**Figure 1:**
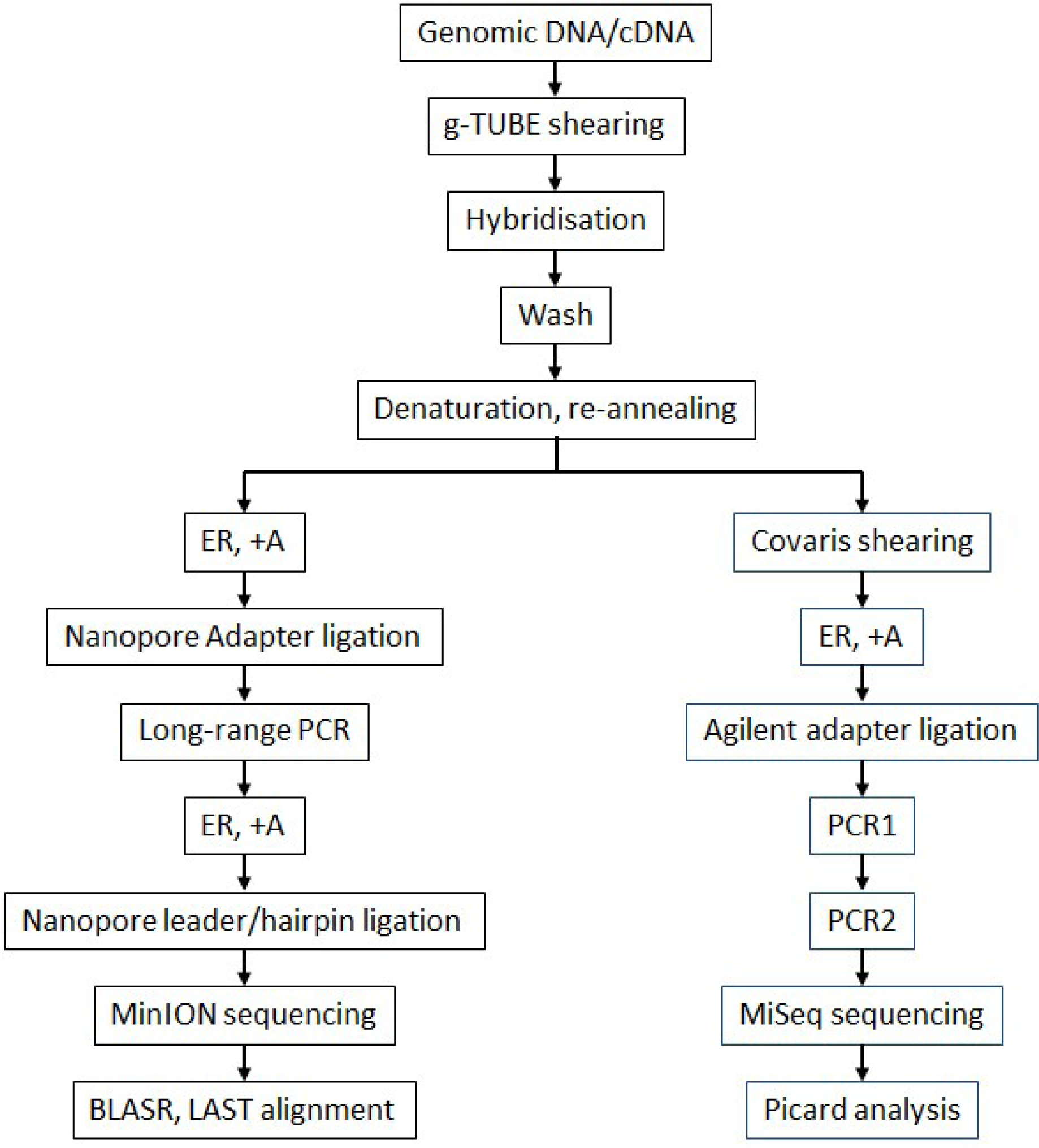
Workflow for hybridisation and sequencing of long-fragment-enriched pathogen DNA. ER: end repair of fragments, *A: dA-tailing.

Hybridised samples were split into two aliquots and half was used for Nanopore library preparation with ONT kit versions SQK-MAP003 for *Mtb* H37Rv and SQK-MAP004 for CMV, FluA, *Mtb*C. The remainder was used to generate Illumina-compatible libraries, using another modified version of the SureSelect^XT^ Target Enrichment for Illumina Paired-End Multiplexed Sequencing (Figure 1).

### Nanopore library preparation, sequencing and analysis

Following renaturation, end repair and dA-tailing were performed with enzymes from the SureSelect^XT^ kit (#5500-0075, Agilent) as specified by the manufacturer. AMPure XP-purified, dA-tailed samples were ligated for 15 min at room temperature in 50 μl volume with 20 μl PCR adapters (ONT, sequencing kits SQK-MAP003 and SQK-MAP004), 5 μl 10x T4 DNA ligase reaction buffer and 2 μl T4 DNA ligase (Agilent). They were cleaned up with 90 μl AMPure XP beads and eluted in 50 μl H_2_O. This ligated DNA (48 μl) was amplified by long-range PCR with 2 μl PCR primers (ONT) and 50 μl Long Amp Taq 2x Master mix (#M0287, NEB) with the following program: 95°C 3 min, 15-18 cycles of [95°C 15 sec, 62°C 15 sec, 65°C 10 min], 65°C 20 min, 4°C hold. PCR reactions were cleaned up with 100 μl AMPure XP beads and eluted in 50 μl H_2_O. Concentrations and fragment sizes were measured with a Qubit fluorometer and Agilent Tape Station as before.

A second round of end repair and dA-tailing was performed on 1 μg of enriched, amplified PCR product using SureSelect^XT^ reagents as described above, but without purification after dA-tailing. Instead, leader/hairpin ligation and sample clean-up were performed according to the ONT protocol for kit SQK-MAP003 (for strain *Mtb* H37Rv only) or SQK-MAP004, in protein LoBind tubes (1.5 ml, #0030108116, Eppendorf). In detail, 10 μl adapter mix and 2 μl hairpin adapter (ONT) were incubated for 10 min at room temperature with 30 μl dA-tailed sample and 50 μl 2x blunt/TA ligase master mix (#M0367, NEB) in 100 μl volume. SQK-MAP003 libraries were cleaned up with AMPure XP beads, SQK-MAP004 with Dynabeads for His-Tag isolation and pulldown (#10103D, Life Technologies) according to the respective ONT protocols. Libraries were eluted from the beads at room temperature for 10 min with 25 μl elution buffer (ONT). Library concentrations were typically 2-10 ng/μl, as assessed by Qubit fluorometer.

Before each MinION^TM^ run, flowcells were quality-tested with the script MAP_Platform_QC (MinKnow software version 0.46.2.8 to 0.49.2.9), then loaded with 6 μl library and 4 μl fuel mix in 140 μl EP buffer (ONT), and run with script MAP_48Hr_Sequencing_Run, typically for 24h. Reads were analysed by the Metrichor 2D basecalling (versions 2.19 to 2.29) cloud-based platform, and the resulting fast5 files (“pass” and “fail” quality) converted to fasta format with Poretools^8^. BLASR^9^ and LAST^10^ were used to align reads to the pathogen reference sequences (human CMV herpes virus HHV-5 GU179001.1, *Mtb* strain H37Rv NC_018143.2, and Influenza strain H1N1, A/Puerto Rico/8/1934). Files were further tested with both aligners against background human (Human_g1k_v37, www.1000genomes.org) or dog (Ensembl CanFam3.1 GCA_000002285.2; NC_006583.3) sequences, and the ONT adapters used for PCR.

The FluA control sample that did not undergo hybridisation (75 ng) was PCR-amplified to 500 ng as described previously, then prepared as recommended in the ONT Genomic DNA sequencing protocol SQK-MAP004. For the non-hybridised CMV sample, 500 ng was used directly for Nanopore library preparation (SQK-MAP004).

### Illumina library preparation from long, hybridisation-enriched fragments

After the long-fragment hybridisation, samples of *Mtb* H37Rv, *Mtb*C, CMV and FluA were sheared with a Covaris AFA instrument (Covaris, Woburn MA, USA) to 200 nt fragment size and converted into Illumina-compatible libraries using a modified SureSelect^XT^ protocol (Figure 1) and Agilent reagents as before. Briefly, samples were end-repaired, dA-tailed, adapters ligated, and DNA amplified (6 cycles) as described in the protocol. Following sample purification, the PCR products were re-amplified using post-capture indexed PCR primers for a further 15 cycles. The resulting indexed libraries were quantified by Qubit and Agilent Tape Station as before, and pooled. Sequencing was performed on an Illumina MiSeq machine with paired-end 600V3 kits (#MS-102- 3003) with automatic adapter trimming. Results from the Illumina MiSeq runs were analysed with the Picard pipeline (http://broadinstitute.github.io/picard/).

## Results

### Comparison of Nanopore library size and read length

CMV and *Mtb* gDNA was sheared to fragments of 10-15 kb using Covaris g-TUBEs. FluA cDNA was used without previous shearing for both enriched and non-enriched Nanopore libraries. Table 1 shows the median fragment length of the input DNA after shearing, as determined on an Agilent Tape Station. Interestingly, the distribution of the unsheared FluA cDNA on the Tape Station showed distinct peaks of 120 nt, 300 nt, 500 nt, 800 nt, 1-3 kb, with fragments up to 15 kb; these correspond to transcripts from the eight FluA genomic segments NC_002016 to NC_002023, and cell line DNA. PCR-amplified samples had wide ranges of sizes both within and between individual reactions, with PCR products about half the size of the original DNA used for hybridisation. The Nanopore reads (raw data in the European Nucleotide Archive, Study PRJEB12651) were similarly variable in length, as indicated by the size of the standard deviations in Table 1. “Pass” quality (both strands read while passing through the nanopore, resulting in higher confidence) reads were, on average, 1.2 to 4.2x the length of “fail” quality reads. PCR-amplified samples were considerably smaller than the original DNA fragments. Sequenced reads were shorter still, but with a wide range, reflected in high standard deviations in Table 1. Reads from non-hybridised samples were longer than enriched samples, either due to fragment damage during the hybridisation and wash processes, or during PCR, which preferentially amplifies shorter fragments.

**Table 1:** Average DNA fragment sizes. The table shows input, post-PCR library size, and average Nanopore read length (“pass” and “fail” quality) with standard deviations (SD) of the samples used in this study. The non-hybridised CMV sample was not PCR-amplified as enough material was available to proceed directly to sequencing. *FluA samples were not sheared.

| Samples/input sizes | Sheared (nt) | PCR-amplified (nt) | Sequenced pass reads (nt/nt SD) | Sequenced fail reads (nt/nt SD) |
| --- | --- | --- | --- | --- |
| FluA non-hybridised | 120, 300, 500, 800, 1000, 3000+* | 99-4000 | 1598/1191 | 805/946 |
| FluA | 120, 300, 500, 800, 1000, 3000+* | 370-4000 | 773/683 | 533/733 |
| CMV non-hybridised | 1960, 49000 | - | 3176/2291 | 487/1203 |
| CMV | 12500 | 1587, 5640 | 1528/975 | 1083/1099 |
| *Mtb* H37Rv | 13800 | 2000-7000 | 2402/1865 | 757/1855 |
| *Mtb*C | 15000 | 1500 | 759/355 | 596/713 |

### Comparison of BLASR and LAST aligners

We used BLASR^9^ and LAST^10^ (with the settings used in Quick et al.^11^) for the alignment of Nanopore reads to their respective references (pathogen and human/dog cell line). We found that BLASR alignment resulted in fewer reads, with higher identity to the reference strains, and lower standard deviation. In contrast, the LAST aligner produced more reads aligning to the reference, with lower identity and higher standard deviation. Table 2 shows statistics for the similarities to the target references obtained with the two aligners.

**Table 2:** Average similarity and length (with standard deviations, SD) of Nanopore reads aligned to the pathogen targets using BLASR and LAST.

| Sample | BLASR alignment |  | LAST alignment |  |
| --- | --- | --- | --- | --- |
|  | Similarity of reads to target (%/SD %) | Length of reads (nt/SD nt) | Similarity of reads to target (%/SD %) | Length of reads (nt/SD nt) |
| **FluA hybridised** | 80/6.3 | 185/109 | 74.9/6.9 | 344/150 |
| **CMV hybridised** | 79.6/6.4 | 940/835 | 72.1/7.9 | 1576/1102 |
| ***MtbC* hybridised** | 80.3/6 | 305/224 | 71.8/8.1 | 628/958 |
| ***Mtb* H37Rv hybridised** | 77.9/5.7 | 1182/1056 | 71.9/8.3 | 1744/1255 |

Most “pass” quality reads aligned to either the target organism or the respective cell line, whereas most “fail” quality reads did not match to either target, cell line, or sequences in the PubMed Nucleotide database (November 2015). A small number of FluA (85), CMV (315) and *Mtb*C (10) Nanopore reads matched to both target pathogen and cell line, predominantly in the output of the LAST aligner (Table 3). However, when compared to the Nucleotide database (http://blast.ncbi.nlm.nih.gov/Blast.cgi, November 2015), these reads only aligned to either reference and not both, indicating suboptimal performance of these aligners for Nanopore reads.

**Table 3:**
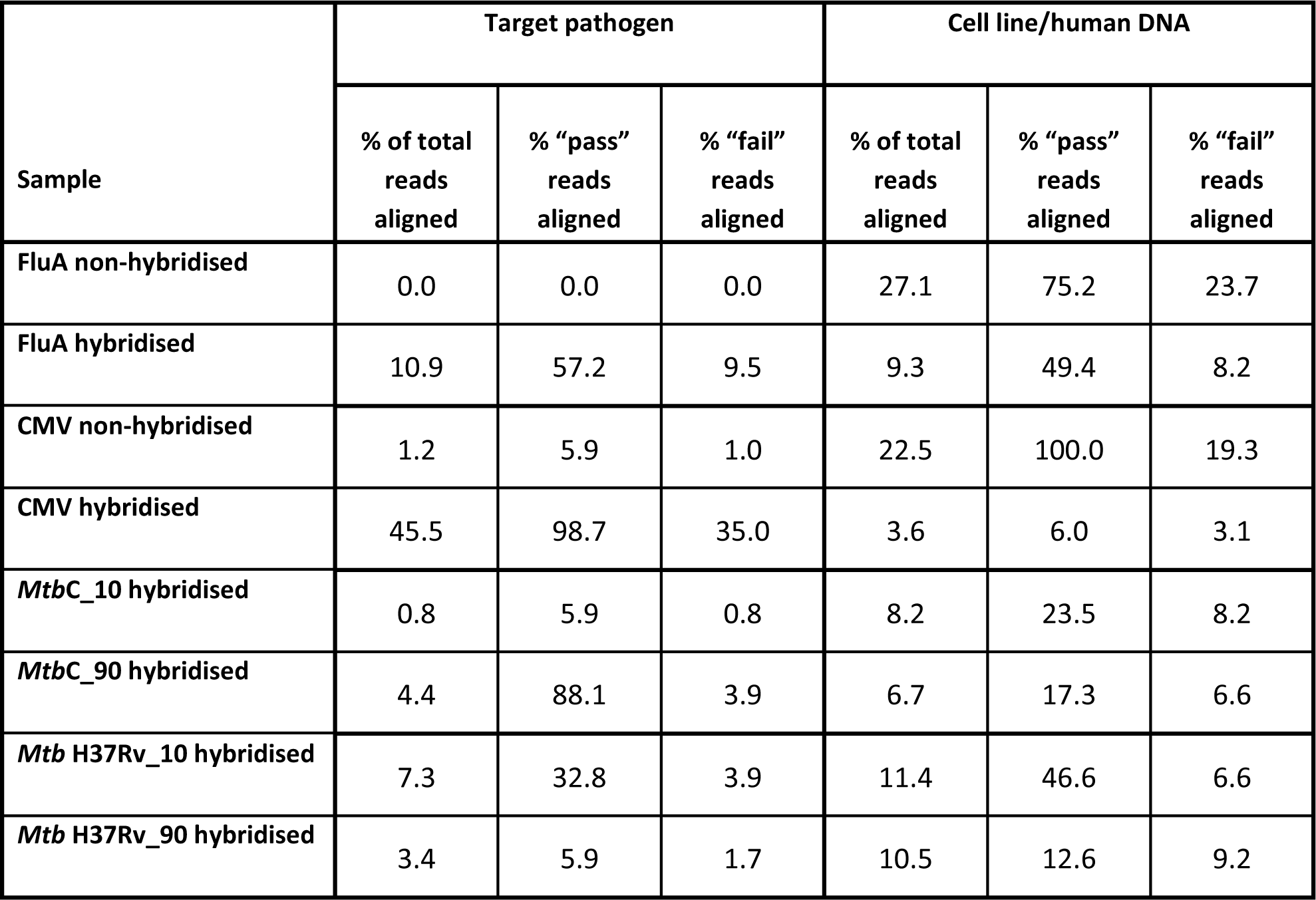
Percentages of reads aligned to target pathogen and cell line/human DNA in the samples prepared for this study.

### Comparison of enriched and non-enriched Nanopore libraries

Amplification and subsequent sequencing of the long DNA fragments demonstrates the success of the hybridisation and library preparation for both Nanopore and Illumina sequencing. Analysis of the 42261 reads obtained from a non-enriched, PCR-amplified FluA cDNA library run on the Nanopore MinION^TM^ found 75.2% pass and 23.7% fail reads aligned to the MDCK dog cell line used for cultivation of the virus, whilst only one read aligned to the FluA reference H1N1. After hybridisation and amplification, 57.2% of pass and 9.5% of fail reads (34211 reads in total) from a Nanopore run could be aligned to FluA. This amounts to a total FluA coverage of 71.2x. Though there was generally good coverage across the eight FluA segments, Figure 2 shows a wide variation of number of reads per fragment, as well as distinct peaks of coverage of FluA cDNA. Similarly, the frequency of cell line reads dropped to 49.4% (pass) and 8.2% (fail) (Table 3).

**Figure 2:**
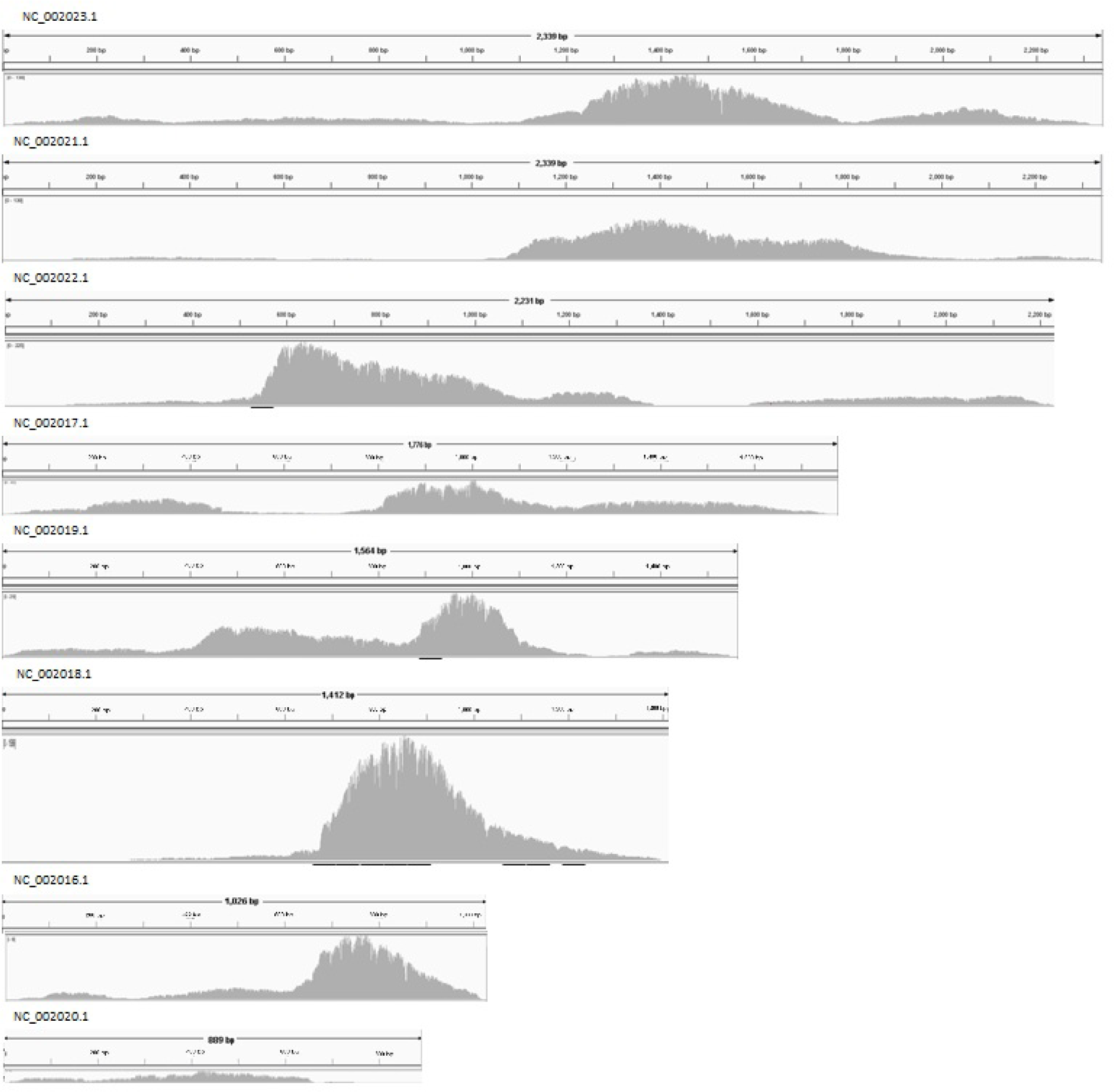
Nanopore reads from enriched FluA cDNA, aligned to reference FluA H1N1 with BLASR, visualized in the Integrated Genome Viewer^12, 13^ (IGV) Maximum coverage results for the fragments are: 139 (NC_002023.1), 139 (NC_002021.1), 225 (NC_002022.1), 51 (NC_002017.1), 219 (NC_002019.1), 1589 (NC_002018.1), 185 (NC_002016.1), 16 (NC_002020.1).

The un-enriched CMV library (432 unique reads in total) produced five reads (1.2%) matching the CMV reference HHV-5, while 97 reads (22.5%) matched the human_g1k_v37 reference. After hybridisation of the DNA with the CMV-specific bait set, we obtained 37589 reads from three runs, with almost all (98.7%) pass reads and 35% of fail reads aligning to the CMV reference. This amounts to a coverage of 89.9x of the CMV genome. Figure 3 shows the output of these reads aligned to the reference.

**Figure 3:**
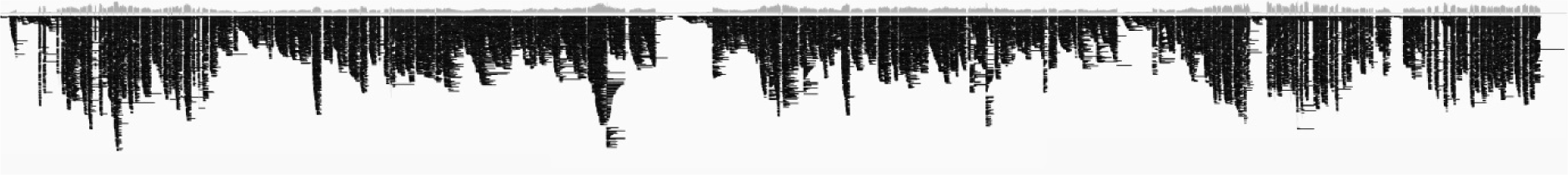
Coverage of the CMV strain HHV-5 genome with Nanopore reads, aligned with BLASR and visualized with IGV. The maximum coverage for this plot is 291.

A comparison of the consensus sequence generated from the enriched CMV reads aligned to the CMV HHV-5 reference using the genomic similarity search tool YASS^14^ found the former had 99.4% similarity to the reference (233854 of 235230 nucleotide residues). The conflicting/mismatch residues are mostly gaps in the Nanopore consensus sequence at 46364-46433 (proteins UL34 and UL35), 147820-147830 (helicase-primase subunit UL102), 194363-194698 and 195851-95977. The last two regions of difference coincide with inverted repeat regions^6^ (194344 to 195667, 195090 to 197626). A number of mismatches to the reference HHV-5 were identified upstream of base 1270; these were due to low coverage of this region by Nanopore reads. We found regions with low (<5x) coverage had a high number of mismatches compared to the reference, but areas of greater coverage matched near-perfectly.

For *Mtb* strain H37Rv, we obtained 2028 unique pass and 9961 unique fail reads (0.077x coverage), for the strain *Mtb*C, 202 pass and 46711 fail reads (0.182x). Distribution of reads from both *Mtb* strains aligning to the H37Rv genome is relatively even (Figure 4). Localized high-coverage areas with multiple reads in both strains were found at a number of genomic positions encoding transposases, e.g. 887488-887429, 890363-889044, 1539822-1538580, 2640242-2635594, 3547252-3544391, 3789669-3788312 of strain H37RV (NC_018143.2). The areas with increased coverage can also be observed in Illumina-generated datasets (Figure 4), presumably due to the redundancy of the sequence, which could result in localised increased aligning of reads.

**Figure 4:**
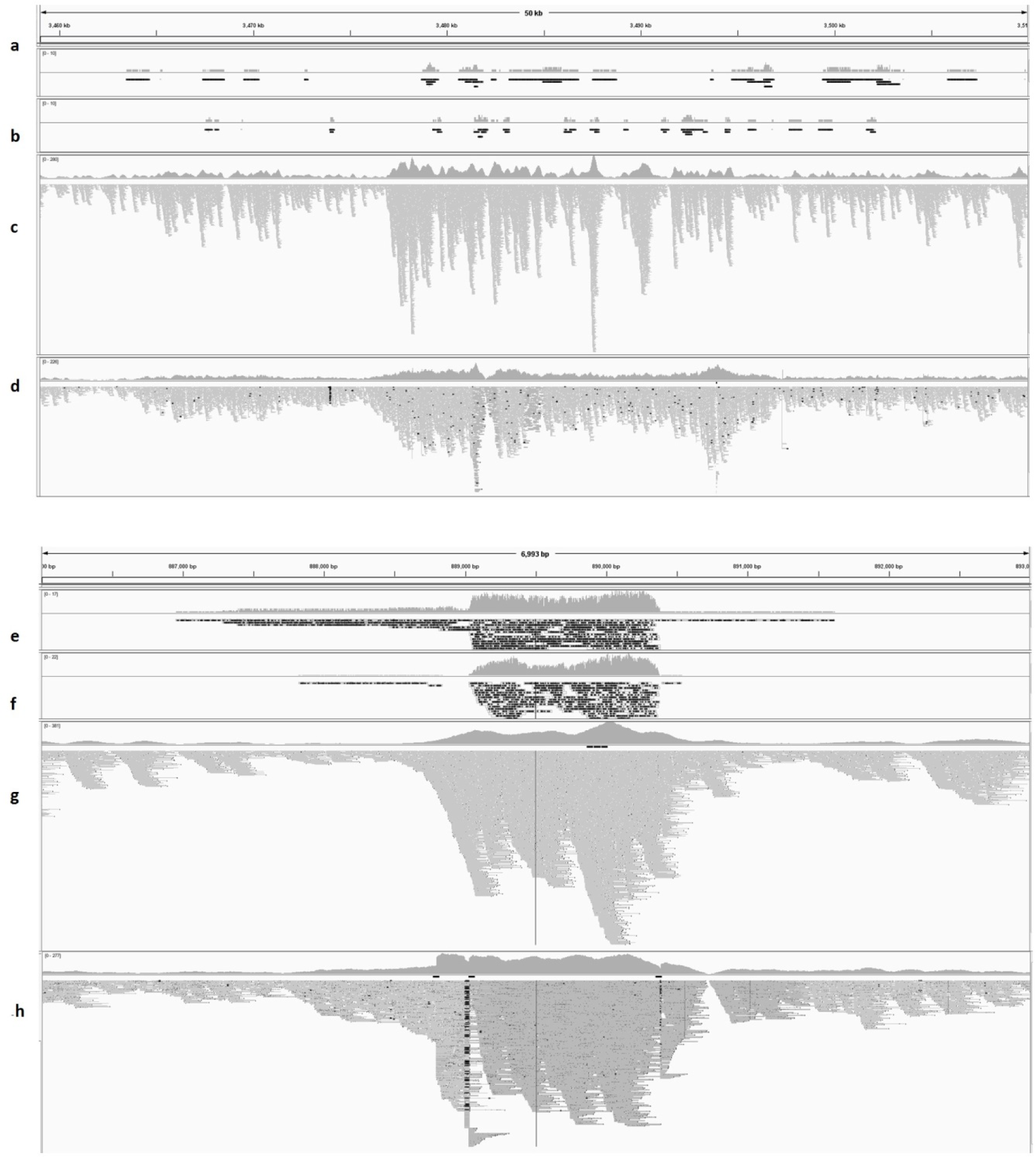
Alignments of Nanopore reads of libraries from DNA of strains *Mtb* H37Rv (a), *Mtb*C (b) and Illumina-sequenced long-fragment-hybridised *Mtb* H37Rv (c) and *Mtb*C (d), in position 3,459,000-3,510,000 of the genome of strain H37Rv show low but even coverage by Nanopore reads (a-d). Panels e-h show the same samples in the same order, in a region of high coverage (886,000-893,000), visualised in IGV. Regions with increased coverage in the Nanopore reads correspond to higher read depth in the Illumina coverage.

### Sequencing of enriched long fragments on the Illumina MiSeq

To assess the success of the long fragment hybridisation, Illumina libraries were generated from half of the hybridised material, and sequenced on a MiSeq (Table 4). The coverage profile from the enriched FluA and *Mtb* samples produced similar distribution, and reads from both the Nanopore and Illumina sequencers showed preferential enrichment of the same regions (Figure 4 shows *Mtb*). In contrast, aligned Nanopore CMV reads generated from the long fragment-enriched sample had a slightly different coverage profile to their Illumina counterpart. The longer Nanopore reads formed wide peaks, whilst mapped Illumina reads clustered in narrow stacks with deep troughs (Figure 5). This could be due to the positioning of baits, which could preferentially enrich short fragments that overlap well with them. Some gaps are visible in the Nanopore coverage, these are bridged by relatively few reads and do not always correspond to gaps in the Illumina coverage.

**Figure 5:**
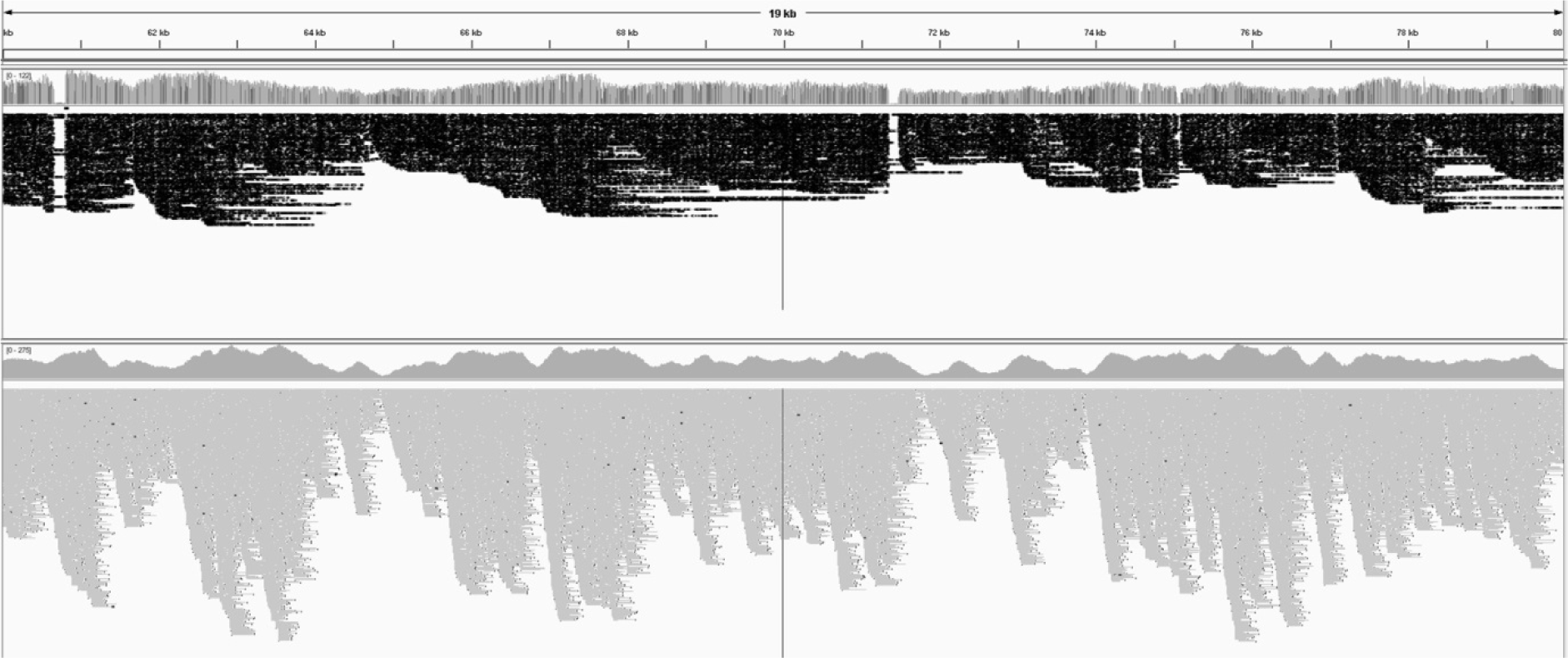
Alignment of Nanopore reads (top) and Illumina reads (bottom) to reference human CMV herpes virus HHV-5 GU179001.1, positions 60,000-80,000, visualised in IGV. The longer Nanopore reads cover the reference more evenly but show some gaps that are well-covered by Illumina reads.

**Table 4:** Statistics of coverage generated by Picard (http://broadinstitute.github.io/picard/) in Illumina MiSeq runs of independent hybridisations of long fragments.

| Sample | Fragment hybridisation | Number of reads aligned to target pathogen | % reads aligned to target pathogen | Mean depth of pathogen coverage | % target bases covered at 10x |
| --- | --- | --- | --- | --- | --- |
| FluA | Long | 2664967 | 59 | 2089 | 97 |
| CMV | Long | 1765332 | 94 | 957 | 96 |
| 10% *Mtb* H37Rv (1) | Long | 6906339 | 84 | 295 | 99 |
| **10% Mtb H37Rv (2)** | Long | 1071332 | 62 | 50 | 99 |
| **10% Mtb H37Rv (3)** | Long | 8193188 | 96 | 315 | 99 |
| **10% Mtb H37Rv (4)** | Long | 2942898 | 56 | 100 | 98 |
| **90% Mtb H37Rv (5)** | Long | 2258868 | 93 | 99 | 99 |
| **90% Mtb H37Rv (6)** | Long | 6978112 | 97 | 297 | 99 |
| **90% Mtb H37Rv (7)** | Long | 9926437 | 87 | 342 | 99 |
| **90% Mtb H37Rv (8)** | Long | 15382452 | 96 | 521 | 98 |
| **Mtb H37Rv 9)** | Short | 3982148 | 99 | 169 | 99 |
| **90% MtbC (1)** | Long | 689141 | 18 | 24 | 85 |
| **MtbC (2)** | Short | 2980023 | 99 | 115 | 97 |

## Discussion

This study explores the enrichment of long fragments using baits designed and used in previous studies^4, 5^ for whole-genome sequencing of specific pathogens from clinical or mixed samples. These, instead of the 200 nt strands conventionally used for Illumina library construction, give a wider range of possibilities for the deconvolution of repeat regions, detection of translocations and larger indels, or mate-pair libraries. Pathogens with small genomes, in our case cDNA from FluA, could be sequenced without previous shearing, potentially preserving structural information and avoiding assembly problems. We show that the enriched long fragments can be used for libraries for both Illumina sequencers (after re-shearing), and the third-generation long-range sequencer Oxford Nanopore MinION^TM^. The portable, relatively inexpensive, and low-footprint MinION^TM^ sequencers have been used in settings where conventional Illumina sequencing would be difficult^15^. Nanopore sequencing has previously been used to detect structural variants in large genomes^16^; our method could be employed as a non-amplicon-based alternative for this application. As the enrichment approach is platform-agnostic, it could also be used to generate libraries compatible with the PacBio RS II and Sequel systems.

Previous work^17^ has shown that detection of moderate to high titres of pathogen DNA (chikungunya virus, Ebola and Hepatitis C virus) from human blood samples is possible using Nanopore sequencing. However, this direct sequencing approach cannot take full advantage of the capabilities of sequencing for strain typing and variant identification, especially in bacterial infections, due to their larger genomes, and/or low titre samples. In our Nanopore sequencing runs with un-enriched FluA or CMV DNA, we detected very low numbers of reads from the pathogen compared to those from the host cell line culture. In contrast, sequencing data from enriched DNA produced good coverage of the FluA and CMV genomes and partial coverage of the *Mtb* genome. High and even coverage of the target pathogen genome^18^, gained through targeted enrichment, can compensate for the errors seen in Nanopore sequencing, and facilitate the detection of potential resistance to antibiotics and identification of mixed infections and minor variants.

We found Illumina sequencing results from material derived from long fragment hybridisation (Table 4) showing good and even coverage, similar to the standard PATHSEEK method of short-fragment hybridisation^4, 5^. This, together with the fact that high-molecular weight post-enrichment fragments could be readily ligated and amplified, highlights the suitability for long fragments for hybridisation and its downstream applications.

We analysed our Nanopore datasets with two different aligners - BLASR produced fewer reads with higher identity to the target pathogen and lower standard deviation, LAST identified more reads with lower identity and higher standard deviation, similar to the results of Kilianski et al.^19^. The difference of up to 40% in sequences identified as matches to the reference between the two aligners, highlights that neither works optimally for aligning Nanopore reads to their reference. A small number of reads reported as containing both cell line and pathogen matches, mainly from the output of LAST, could not be confirmed. As reported elsewhere^17, 19^, a large percentage of Nanopore reads (mainly the “fail” quality) could not be aligned to either the target pathogen, or human (*Mtb*), the human cell line (CMV) or the dog cell line (FluA), respectively, and show no similarities when compared to the NCBI Nucleotide Database (November 2015).

The persisting, and, in some cases, increasing pressure of pathogenic viruses and bacteria on human and animal health underlines the need for fast, accurate and up-to-date diagnosis. Next-generation sequencing can enable clinicians to identify both pathogen species and genotype in one step, significantly aiding individual treatment with respect to drug resistance^20^ and monitoring of outbreaks^21^. The increasing interest in whole-genome sequencing for diagnosis has led to the drive to take it out of the research environment and into the clinic to reduce the time between taking a specimen, identifying the pathogen and obtaining actionable genomic data. Nanopore sequencing, coupled with data streaming and real-time analysis, has the possibility to bring sequencing closer to patient, or to be used in settings without easy access to an Illumina machine. The Nanopore sequencing platform, coupled with our long-fragment enrichment method, can be applied to a range of pathogens and clinical samples. However, our experiments were performed with a 16-hour hybridisation reaction, a time-consuming and rate-limiting step. In the future, this has the potential to be shortened to four hours by using a different hybridisation protocol. Though not tested in this study, addition of molecular bar-codes as outlined in the “Sequencing using the PCR Barcoding Kit” ONT protocol, will allow for several clinical samples of enriched viral DNA to be run simultaneously on one MinION^TM^ flowcell. This, coupled with increasing accuracy of the MinION^TM^ reads, will also reduce the coverage necessary for strain and variant identification, making this method suitable for diagnostic purposes in the future.

## Acknowledgements

We would like to thank Dietrich Lueersen, David Blaney and Dan Swan for their help in the analysis of the data; Richard Milne at Department for Virology, UCL Medical School (Royal Free Campus, Rowland Hill Street, London, UK) for the kind gift of CMV Merlin strain, Amanda Brown, Lise J Schreuder and Tanya Parish at the Barts and The London School of Medicine and Dentistry (Queen Mary University of London, London, UK), for strain H37Rv, Philip Butcher *Mtb* H37Rv, Philip Butcher and Jasvir Dhillon at St. George’s Hospital (University of London, London, UK) for the generous donation of strain *Mtb* C. Past and present members of the PATHSEEK consortium are: Judith Breuer, Rachel Williams, Mette Theilgaard Christiansen, Josie Bryant, Sofia Morfopoulou, Helena Tutill, Erika Yara-Romero, Charlotte Williams, Dan Depledge (UCL), Martin Schutten, Saskia Smits, Georges M.G.M. Verjans, Freek B. van Loenen, Anne van der Linden, Albert Osterhaus (Erasmus MC); Katja Einer-Jensen, Martin Ludvigsen, Roald Forsberg (CLC Bio); James Clough, Graham Speight, Jacqueline Chan, Jolyon Holdstock, Sabine Eckert, Mike McAndrew, Amanda Brown (OGT).

## References

1. Carlet J. The world alliance against antibiotic resistance: consensus for a declaration. Clin Infect Dis. 2015; 60(12): 1837–1841.

2. Wlodarska M, Johnston JC, Gardy JL, Tang P. A microbiological revolution meets an ancient disease: improving the management of tuberculosis with genomics. Clin Microbiol Rev. 2015; 28(2): 523–539.

3. Doughty EL, Sergeant MJ, Adetifa I, Antonio M, Pallen MJ. Culture-independent detection and characterisation of *Mycobacterium tuberculosis* and *M. africanum* in sputum samples using shotgun metagenomics on a benchtop sequencer. PeerJ. 2014; 2: e585.

4. Christiansen MT, Brown AC, Kundu S, Tutill HJ, Williams R, Brown JR, et al. Whole-genome enrichment and sequencing of Chlamydia trachomatis directly from clinical samples. BMC Infect Dis. 2014; 14: 591.

5. Brown AC, Bryant JM, Einer-Jensen K, Holdstock J, Houniet DT, Chan JZ, et al. Rapid Whole-Genome Sequencing of Mycobacterium tuberculosis Isolates Directly from Clinical Samples. J Clin Microbiol. 2015; 53(7): 2230–2237.

6. Masse MJ, Karlin S, Schachtel GA, Mocarski ES. Human cytomegalovirus origin of DNA replication (oriLyt) resides within a highly complex repetitive region. Proc Natl Acad Sci U S A. 1992; 89(12): 5246–5250.

7. Witney AA, Gould KA, Arnold A, Coleman D, Delgado R, Dhillon J, et al. Clinical application of whole-genome sequencing to inform treatment for multidrug-resistant tuberculosis cases. J Clin Microbiol. 2015; 53: 1473–1483.

8. Loman NJ, Quinlan AR. Poretools: a toolkit for analyzing Nanopore sequence data. Bioinformatics 2014; 23: 3399–3401.

9. Chaisson MJ, Tesler G. Mapping single molecule sequencing reads using basic local alignment with successive refinement (BLASR): application and theory. BMC Bioinformatics 2013; 13: 238.

10. Kielbasa SM, Wan R, Sato K, Horton P, Frith MC. Adaptive seeds tame genomic sequence comparison. Genome Research 2011; 21(3): 487–493.

11. Quick J, Quinlan AR, and Loman NJ. A reference bacterial genome dataset generated on the MinION^TM^ portable single-molecule nanopore sequencer. GigaScience 2014; 3: 22.

12. Robinson JT, Thorvaldsdóttir H, Winckler W, Guttman M, Lander ES, Getz G, et al. Integrative Genomics Viewer. Nature Biotechnology 2011; 29: 24–26.

13. Thorvaldsdóttir H, Robinson JT, Mesirov JP. Integrative Genomics Viewer (IGV): highperformance genomics data visualization and exploration. Briefings in Bioinformatics 2013; 14: 178–192.

14. Noe L, Kucherov G. YASS: enhancing the sensitivity of DNA similarity search. Nucleic Acids Res. 2005; 33(2): W540–W543.

15. Quick J, Loman NJ, Duraffour S, Simpson JT, Severi E, et al. Real-time, portable genome sequencing for Ebola surveillance. Nature 2016; 530(7589):228–32.

16. Norris AL, Workman RE, Fan Y, Eshleman JR, Timp W. Nanopore sequencing detects structural variants in cancer. Cancer Biol Ther. 2016; 17(3):246–53.

17. Greninger AL, Naccache SN, Federman S, Yu G, Mbala P, Bres V, et al. Rapid metagenomic identification of viral pathogens in clinical samples by real-time nanopore sequencing analysis. Genome Med. 2015; 7: 99.

18. Judge K, Harris SR, Reuter S, Parkhill J, Peacock SJ. Early insights into the potential of the Oxford Nanopore MinION for the detection of antimicrobial resistance genes. J Antimicrob Chemother. 2015; 70(10): 2775–2778.

19. Andy Kilianski A, Haas JL, Corriveau EJ, Liem AT, Willis KL, Kadavy DR, et al. Bacterial and viral identification and differentiation by amplicon sequencing on the MinION nanopore sequencer. GigaScience 2015; 4: 12.

20. Köser CU, Ellington MJ, Peacock SJ. Whole-genome sequencing to control antimicrobial resistance. Trends Genet. 2014; 30(9): 401–407.

21. Chin CS, Sorenson J, Harris JB, Robins WP, Charles RC, Jean-Charles RR, et al. The Origin of the Haitian Cholera Outbreak Strain. N Engl J Med. 2011; 364(1): 33–42.

